# The best formula for estimating the low density lipoprotein cholesterol on a North African population

**DOI:** 10.1101/083790

**Authors:** Benghezel Hichem, Boukrous Hanane, Zehaf Mouna, Saadi Radia

## Abstract

We often use the estimation of low density lipoprotein cholesterol by using the Friedwald Formula, however limitation and uncertainties make calculation its limits use. Otherwise, simple and inexpensive formulas exists, rarely used in clinical laboratories: Hatorri, Puavilai, Anandaraja, Ahmadi, Vujovic, Saidullah and Cordova. These formulas were studied on small geographically and ethnically populations and require validation in a larger population.

We intended by this work, highlight the formula that best estimates the low density lipoprotein cholesterol more accurately than the Friedewald formula on a North African population.

It seems that the Puivalai formula is the most suitable to be applied on this population.

## INTRODUCTION

A clear link exists between elevated serum low density lipoprotein cholesterol (LDLc) and cardiovascular risk, hence the need for precise dosing, or at least a proper assessment of serum levels of LDLc.

In addition to the reference method (β quantification)^1^, A great live assay panel is currently available (DAIICHI process, Kyowa process, DENKA SEIKEN process and WAKO process)^2^. For economic reasons, biology medical laboratories often use an estimation of LDLc and using the Friedewald formula^3^.

Usage limits and calculation uncertainties often occur when this equation is used in patients with type III hyperlipidemia (Fredrickson classification) and in the presence of high or low concentrations of triglycerides respectively higher than 400 mg / dl and less than 100 mg / dl^4,5^. Thus, an erroneous estimation is demonstrated in diabetics and patients with chronic renal failure or liver failure^6^.

Several other formulas as simple as Friedewald emerged, we cite those Hattori and al^7^ (1998), Puavilai and al^8^ (2004), Anandaraja and al^9^ (2005), Ahmadi and al^5^ (2008), Vujovic and al^10^ (2010), Saiedullah and al^11^ (2009) and Cordova and al (2013)^12^. However, these formulas were studied on small geographically and ethnically populations and require validation in a larger population.(Table 1)

**Table 1:** The formulas of calculating the LDLc

| Name | Formulas |
| --- | --- |
| Friedewald and al | $TC - (HDLc + TG/5)$ |
| Hattori and al | $0,94TC - 0,94HDLc - 0,19TG$ |
| Puavilai and al | $TC - (HDLc + TG/6)$ |
| Anandaraja and al | $0,9TC - 0,9(TG/5) - 28$ |
| Ahmadi and al | $(TC/1,19) + (TG/1,9) - (HDLc/1,1)$ |
| Vujovic and al | $TC - (HDLc + TG/6,85)$ |
| Saiedullah and al | $TC - (HDLc + TG/5) + (15,3TG/(TC - 12,4))$ |

TC: total cholesterol, TG: triglycerides, HDLc: high density lipoprotein cholesterol
LDLc: Low density lipoprotein cholesterol.

We intended by this work, highlight the formula that best estimates the LDLc more accurately than the Friedewald formula on a North African population (Algeria).

## MATERIALS AND METHODS

This retrospective study is performed on 725 serum samples analyzed in the Biochemistry laboratory of the hospital in Batna, Algeria.

The samples were taken after 12 hours of fasting, dry tube and all samples were analyzed in the first hours of their arrival at the laboratory. After clotting at room temperature, serum is separated by centrifugation at 3500 g for 10 min.

We eliminated from our study and systematically samples with the level of triglycerides greater than 1500 mg / dl and hemolytic or icteric serum to avoid the analytical interference to determination of LDLc level.

The determination of cholesterol, triglycerides, high density lipoprotein cholesterol (HDLc) and LDLc was carried out according to the principles of available assay on COBAS INTEGRA 400, Roche Diagnostics. The calibrating and internal controls are provided by the Roche diagnostics company.

LDLc is assayed by the method of DAIICHI, this method has a detection limit estimated at 0.39 mg / dl, repeatability and good reproducibility (CV <5%) with a strong correlation with the reference method 0.954 (Roche diagnostics cobas integra 400/700/800 LDL-D 10/2000, version 1.0).

All values are expressed as mean ± standard deviation. The t test was used to compare means of various formulas with the LDLc assay method and the Pearson correlation test is performed to examine the various correlations. P values ≤ 0.05 were considered statistically significant. The IBM®SPSS 20.0 statistics software to evaluate all donated.

## RESULTS

For 725 samples, the average concentrations of total cholesterol, triglycerides, hight density lipoprotein cholesterol (HDLc) and assayed LDLc are respectively 176 +/− 47mg/dl, 135 +/− 64 mg/dl, 45 +/− 23mg/dl and 108 +/− 42 mg / dl.

A strong correlation was observed between the measured LDLc and other formulas. But, when comparing mean, only the Puavikai et al formula shows no significant difference with LDLc measured. (Table 2)

**Table 2:** Comparison of different formulas with the measured LDL

|  |  | Correlation | P value | t test | P value |
| --- | --- | --- | --- | --- | --- |
| Pair 1 | LDL & Friedwald and al | ,930 | ,000 | 8,2 | ,000 |
| Pair 2 | LDL & Cordova and al | ,919 | ,000 | 16,6 | ,000 |
| Pair 3 | LDL & Vujovic and al | ,934 | ,000 | -4,2 | ,000 |
| Pair 4 | LDL & Ahmadi and al | ,728 | ,000 | -42,8 | ,000 |
| Pair 5 | LDL & Saiedullah and al | ,925 | ,000 | 8,5 | ,000 |
| Pair 6 | LDL & Anandaraja and al | ,840 | ,000 | -5,3 | ,000 |
| Pair 7 | LDL & Puavilai and al | ,933 | ,000 | 581 | ,561 |
| Pair 8 | LDL & Hattori and al | ,930 | ,000 | 19,7 | ,000 |

Secondly we have classified our samples based on the triglycerides level in 04 groups: group 1 (268 samples): triglyceride ≤100 mg / dl (total cholesterol = 152 +/− 39 mg/dl, triglyceride= 77 +/− 24 mg/dl, HDLc= 46 +/− 15 mg/dl and LDLc measured = 95 + −34 mg / dl). The group 2 (316 samples): triglycerides] 100 to 200 [mg / dl (total cholesterol = 180 +/− 44 mg/dl, triglyceride = 138 + / − 27 mg/dl, HDL cholesterol = 46 +/− 26 mg/dl and LDLc = 110 +/− 42 mg / dl). Group 3 (72 samples): triglyceride [200 and 300] mg / dl (total cholesterol = 206 +/− 55 mg/dl, triglyceride= 238 +/− 40 mg/dl, HDLc = 40 +/− 28 mg/dl and LDL cholesterol = 127 +/− 50 mg / dl) and finally, group 4 (28 samples): triglyceride] 300-679] mg / dl ( total cholesterol = 226 +/− 62 mg/dl, triglyceride 283 +/− 85 mg/dl, HDLc= 45 +/− 33 mg/dl and LDLc = 116 +/− 71 mg / dl). Correlations and t test application to different groups gives us the following results. (Table 3)

**Table 3:** Comparison of different formulas with the measured LDL in the four groups

| Group 1 |  | Correlation | *P value* | *t test* | *P value* |
| --- | --- | --- | --- | --- | --- |
| Pair 1 | LDL & Friedwald and al | ,969 | ,000 | 6,739 | ,000 |
| Pair 2 | LDL & Cordova and al | ,963 | ,000 | 21,430 | ,000 |
| Pair 3 | LDL & Vujovic and al | ,969 | ,000 | -1,295 | ,196 |
| Pair 4 | LDL & Ahmadi and al | ,860 | ,000 | -28,822 | ,000 |
| Pair 5 | LDL & Saiedullah and al | ,966 | ,000 | 14,350 | ,000 |
| Pair 6 | LDL & Anandaraja and al | ,903 | ,000 | 10,772 | ,000 |
| Pair 7 | LDL & Puavilai and al | ,969 | ,000 | 1,784 | ,076 |
| Pair 8 | LDL & Hattori and al | ,969 | ,000 | 17,629 | ,000 |
| Group 2 |  | Correlation | *P value* | *t test* | *P value* |
| Pair 1 | LDL & Friedwald and al | ,930 | ,000 | 4,271 | ,000 |
| Pair 2 | LDL & Cordova and al | ,928 | ,000 | 10,722 | ,000 |
| Pair 3 | LDL & Vujovic and al | ,931 | ,000 | -3,872 | ,000 |
| Pair 4 | LDL & Ahmadi and al | ,846 | ,000 | -53,235 | ,000 |
| Pair 5 | LDL & Saiedullah and al | ,927 | ,000 | 4,317 | ,000 |
| Pair 6 | LDL & Anandaraja and al | ,797 | ,000 | -7,099 | ,000 |
| Pair 7 | LDL & Puavilai and al | ,931 | ,000 | -,751 | ,453 |
| Pair 8 | LDL & Hattori and al | ,930 | ,000 | 12,064 | ,000 |
| Group 3 |  | Correlation | *P value* | *t test* | *P value* |
| Pair 1 | LDL & Friedwald and al | ,893 | ,000 | 3,448 | ,001 |
| Pair 2 | LDL & Cordova and al | ,896 | ,000 | 1,211 | ,230 |
| Pair 3 | LDL & Vujovic and al | ,896 | ,000 | -1,264 | ,210 |
| Pair 4 | LDL & Ahmadi and al | ,796 | ,000 | -34,610 | ,000 |
| Pair 5 | LDL & Saiedullah and al | ,872 | ,000 | ,997 | ,322 |
| Pair 6 | LDL & Anandaraja and al | ,894 | ,000 | -8,501 | ,000 |
| Pair 7 | LDL & Puavilai and al | ,895 | ,000 | ,553 | ,582 |
| Pair 8 | LDL & Hattori and al | ,893 | ,000 | 6,355 | ,000 |
| Group 4 |  | Correlation | *P value* | *t test* | *P value* |
| Pair 1 | LDL & Friedwald and al | ,977 | ,000 | 4,240 | ,000 |
| Pair 2 | LDL & Cordova and al | ,974 | ,000 | -4,484 | ,000 |
| Pair 3 | LDL & Vujovic and al | ,982 | ,000 | -3,377 | ,002 |
| Pair 4 | LDL & Ahmadi and al | ,746 | ,000 | -22,388 | ,000 |
| Pair 5 | LDL & Saiedullah and al | ,983 | ,000 | -1,102 | ,280 |
| Pair 6 | LDL & Anandaraja and al | ,854 | ,000 | -7,772 | ,000 |
| Pair 7 | LDL & Puavilai and al | ,981 | ,000 | -,257 | ,799 |
| Pair 8 | LDL & Hattori and al | ,977 | ,000 | 6,512 | ,000 |

We classified each group into subgroups (A, B, C, D) in the functions of total cholesterol levels (≤120 mg / dl] 120-150] mg / dl] 150-200] mg / dl and > 200mg / dl. The set of formulas in different subgroups are highly correlated to the direct assay. When comparing the averages of the different formulas with LDLc measured we obtained this results: In subgroup 1A (58 samples) formulas Puavilai and Vujovic is interchangeable with LDLc measured with respectively p = 0.685 and 0.127 for the 1B sub-group (68 samples) Vujovic seems to be the best formula to estimate LDLc p = 0, 126, for the subgroup 1C (59 samples) formulas Puavilai and Vujovic and are switchable with the assay method with respectively p = 0.628 and p=0.190. Finally, for the 1D sub-group (28 samples) where the formula Friedwald, Anandaraja and Puivilai have p> 0.05 (0.710, 0.378 and 0.06)

For the subgroup 2A (31 samples) formulas Puavilai, Anandaraja, Vujovic Saiedullah, Friedewald and Cordova respectively have a p = 0.88, p=0,952, p=0,445, p=0,235, p=0,180, p=0,91. So as for the subgroup 2B (39 samples) Vujovic and Anandaraja formulas are switchable with the Direct method with p = 0.159 and 0.099. For the subgroup 2C (149 samples) Puavilai is the single best formula with p = 0.385. While for the 2D sub-group (97 samples) Saiedullah and Friedewald formulas are switchable with the measured LDLc (p is respectively equal to 0.616, 0.427).

For group 3 only subgroups C (40 samples) and D (28 samples) are operable with a preference for Vujovic, Puavilai and Saidullah formulas (p = 0.610, 0.590 and 0.105) for C and Puavilai, Saidullah for the D (p = 0.799 and 0.280).

For Group 4 only the subgroup C is most to least important (20 samples) and the formulas for Puavilai and Saidullah each have a p = 0.522 and 0.266. We tried to determine the effect of triglyceride report / cholesterol on choosing the best formula interchangeable with direct assay method, the following results were obtained in table 4.

**Table 4.**

|  | Pair | Correlation | P Correlation | t test | P t test |
| --- | --- | --- | --- | --- | --- |
| ratio < 0,5<br>159 samples | LDL & F | ,965 | ,000 | 3,318 | ,001 |
|  | LDL & C | ,965 | ,000 | 22,741 | ,000 |
|  | LDL & V | ,965 | ,000 | -1,812 | ,072 |
|  | LDL & A | ,952 | ,000 | -24,933 | ,000 |
|  | LDL & S | ,966 | ,000 | 11,435 | ,000 |
|  | LDL & AN | ,844 | ,000 | 3,936 | ,000 |
|  | LDL & P | ,965 | ,000 | ,146 | ,884 |
|  | LDL & H | ,965 | ,000 | 12,415 | ,000 |
| ratio 0,5-1<br>379 samples | LDL & F | ,938 | ,000 | 4,372 | ,000 |
|  | LDL & C | ,939 | ,000 | 14,512 | ,000 |
|  | LDL & V | ,940 | ,000 | -4,123 | ,000 |
|  | LDL & A | ,903 | ,000 | -41,569 | ,000 |
|  | LDL & S | ,937 | ,000 | 6,301 | ,000 |
|  | LDL & AN | ,850 | ,000 | -3,499 | ,001 |
|  | LDL & P | ,939 | ,000 | -,899 | ,369 |
|  | LDL & H | ,937 | ,000 | 13,058 | ,000 |
| ratio 1 -1,5<br>97 samples | LDL & F | ,898 | ,000 | 5,110 | ,000 |
|  | LDL & C | ,918 | ,000 | 1,404 | ,164 |
|  | LDL & V | ,907 | ,000 | -,457 | ,649 |
|  | LDL & A | ,908 | ,000 | -27,991 | ,000 |
|  | LDL & S | ,894 | ,000 | 2,005 | ,048 |
|  | LDL & AN | ,918 | ,000 | -6,018 | ,000 |
|  | LDL & P | ,904 | ,000 | 1,713 | ,090 |
|  | LDL & H | ,898 | ,000 | 8,087 | ,000 |
| ratio 1,5-2<br>39 samples | LDL & F | ,971 | ,000 | 6,913 | ,000 |
|  | LDL & C | ,918 | ,000 | -5,449 | ,000 |
|  | LDL & V | ,963 | ,000 | -2,418 | ,021 |
|  | LDL & A | ,775 | ,000 | -15,901 | ,000 |
|  | LDL & S | ,969 | ,000 | -1,388 | ,173 |
|  | LDL & AN | ,598 | ,000 | -5,294 | ,000 |
|  | LDL & P | ,967 | ,000 | ,796 | ,431 |
|  | LDL & H | ,972 | ,000 | 9,600 | ,000 |
| ratio > 2<br>10 samples | LDL & F | ,816 | ,004 | 2,010 | ,075 |
|  | LDL & C | ,931 | ,000 | -6,708 | ,000 |
|  | LDL & V | ,876 | ,001 | -1,118 | ,293 |
|  | LDL & A | ,896 | ,000 | -11,199 | ,000 |
|  | LDL & S | ,796 | ,006 | -1,677 | ,128 |
|  | LDL & AN | ,936 | ,000 | -6,867 | ,000 |
|  | LDL & P | ,857 | ,002 | ,213 | ,836 |
|  | LDL & H | ,813 | ,004 | 2,558 | ,031 |

## DISCUSSION

We often use the estimation of LDLc using the Friedewald formula, without checking to as the reliability of its results, some as simple and inexpensive formulas that this formula exist, unfortunately they are not widely used in laboratories of biology medical.

The entire population Puavilai formula is largely the best formula for estimating LDLc with a 0.933 correlation coefficient (linear regression y = 0,98x + 0.042 mg / l) and an average of difference with the direct assay method are not statistically significant (Figure 1).

**Figure 1:**
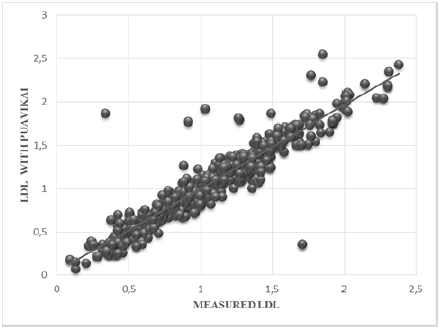
Correlation between LDLc measured and Puavilai formula

It appears from this study that the Friedewald formula often overestimates LDLc and is interchangeable with the direct determination in 20.55% of cases (subgroups 1D, 2A and 2D). The others formulas often comes up is the Vujovic formula, that is switchable with the direct assay method in 40, 68% of cases (subgroup 1A, 1B, 1C, 2A, 2B and 3C). The Saidullah, Ananddaraja and Codova formulas are interchangeable with direct assay in respectively 28.7%, 13.5% and 4.1% of cases.

In the Friedewald formula the ratio LDL-C = TC [HDL-C + TG / 5, the 5 or k indicates the ratio of triglycerides on the cholesterol (5: 1) in the lipoproteins of very low density (VLDL).

A Japanese team Y Haya and al showed a better estimation of LDL cholesterol if k changed with serum triglycerides; K=3 with lower triglycerides 150 mg / dl, 4 for those with triglycerides from 150 to 299 mg / dl, 5 for those with triglycerides of 300 to 400 mg / dl.^13^

We note that the two formulas that emerge most often are identical to the formula of Friedewald with K = 6.85 for Vujovic formula and k = 6 for Puavilai formula. However, in our case the choice of the most switchable formulas do not follow the elevation of triglyceride concentrations. But surprisingly, we note that it follows the ratio of triglyceride total cholesterol, indeed Vujovic formula is only usable when the ratio is between 1 and 1.5 (correlation coefficient 0.907 P <0.0001, t = −0.457 p = 649) but the Puavilai formula against is the more interchangeable with the method of DAIICHI in other situations (correlation coefficient of 0.948 p <0.0001, t = 0.548 p = 0.548).

## CONCLUSION

The Friedewald formula to estimate LDLc seems not to be suitable for the Algerian population. Other formulas include that of Puavilai gives better results, it also appears that the triglyceride cholesterol ratio of in VLDL is closer to 6 than 5.

The development of a formula for calculating the own LDLc our population and applicable to a wide category of patient is of paramount necessity to correctly estimate LDLc is a close as possible to the true LDLc.

